# Structural basis underlying strong interactions between ankyrins and spectrins

**DOI:** 10.1101/2020.03.04.976142

**Authors:** Jianchao Li, Keyu Chen, Ruichi Zhu, Mingjie Zhang

**Author notes:** These authors contributed equally to this work (Jianchao Li, Keyu Chen). Correspondence: Keyu Chen and Mingjie Zhang.

## Abstract

Ankyrins (encoded by *ANK1/2/3* corresponding to Ankyrin-R/B/G or AnkR/B/G), via binding to spectrins, connect plasma membranes with actin cytoskeleton to maintain mechanical strengths and to modulate excitabilities of diverse cells such as neurons, muscle cells, and erythrocytes. Cellular and genetic evidences suggest that each isoform of ankyrins pairs with a specific β-spectrin in discrete subcellular membrane microdomains for distinct functions, though the molecular mechanisms underlying such ankyrin/β-spectrin pairings are unknown. In this study, we discover that a conserved and short extension N-terminal to the ZU5_N_-ZU5_C_-UPA tandem (exZZU) is critical for each ankyrin to bind to β-spectrins with high affinities. Structures of AnkB/G exZZU in complex with spectrin repeats13-15 of β2/β4-spectrins solved here reveal that the extension sequence of exZZU forms an additional β-strand contributing to the structural stability and enhanced affinity of each ZU5_N_/spectrin repeat interaction. The junction site between the extension and ZU5_N_ is exactly the position of a splicing-mediated miniexon insertion site of AnkB/G. The complex structures further reveal that the UPA domain of exZZU directly participates in spectrin binding. Formation of the exZZU supramodule juxtaposes the ZU5_N_ and UPA domains for simultaneous interacting with spectrin repeats 14 and 15. However, our biochemical and structural investigations indicate that the direct and strong interactions between ankyrins and β-spectrins do not appear to determine their pairing specificities. Therefore, there likely exists additional mechanism(s) for modulating functional pairings between ankyrins and β-spectrins in cells.

## Introduction

Ankyrins and spectrins are key scaffold proteins in micron-scale membrane domains of excitable and mechano-resistant tissues/cells, such as the heart T-tubules in cardiomyocytes, the axon initial segment (AIS) and nodes of Ranvier (NOR) in neurons, the lateral membrane in epithelial cells, the sarcomeres in skeletal muscles and the plasma membrane in erythrocytes [1–3]. These specialized membrane domains clustered high density of ion channels, cell adhesion molecules and diverse signaling proteins to achieve their highly specialized functions. By anchoring and stabilizing numerous integral membrane proteins at specific membrane sites, ankyrin and spectrin complexes link these integral membrane proteins to the spectrin/actin based cytoskeletal networks, helping to organize and support these membrane micro-domains. Dysfunctions of ankyrin/spectrin complexes lead to malfunctions of membrane microdomains and cause many different types of human diseases. For example, a number of loss-of-function mutations in ankyrin-R (AnkR) and β1-spectrin are known to cause hereditary spherocytosis [4–6]; genetic variants of ankyrin-G (AnkG) and ankyrin-B (AnkB) are associated with bipolar disorders [7–11], schizophrenia [12–14], autism spectrum disorders [15–22] and severe ataxia [23]; mutations in β3-spectrin cause spinocerebellar ataxia type 5 (SCA5) [24–26]; alterations in β4-spectrin cause spectrinopathy [27–29]; dysfunction of AnkB or β2-spectrin can lead to serious cardiac pathologies, like inherited cardiac arrhythmia that known as the AnkB syndrome [30–37].

Ankyrin family contains three members, AnkR, AnkB and AnkG, which are encoded by *ANK1*, *ANK2* and *ANK*3 respectively. All three ankyrins share similar domain organizations (Figure 1A): a membrane-binding domain (MBD) located in the N-terminus responsible for interacting with various membrane proteins [38], a spectrin-binding domain (SBD) following MBD, a linker region connecting MBD and SBD which is also capable of folding back to inhibit MBD/target interactions [39], an unstructured tail following SBD with extremely different lengths followed by a death domain, and a short C-terminal domain. The SBD is composed of three subdomains, ZU5_N_, ZU5_C_ and UPA domain, forming a structural supramodule (ZZU) [40]. Spectrin family is composed of two groups of subunits: α-spectrins and β-spectrins, each containing multiple continuous spectrin repeats. Under the plasma membrane of living cells, spectrins exist as tetramers, each formed by two α and two β subunits. Vertebrates express two α-spectrins (αI, αII) and five β-spectrins (β1, β2, β3, β4, and β5). Their spectrin repeats endow spectrin tetramers with mechanical resilience [41–44]. Except for β5-spectrin, all mammalian β-spectrins can bind to ankyrins through their middle spectrin repeats [1,45].

**Figure 1:**
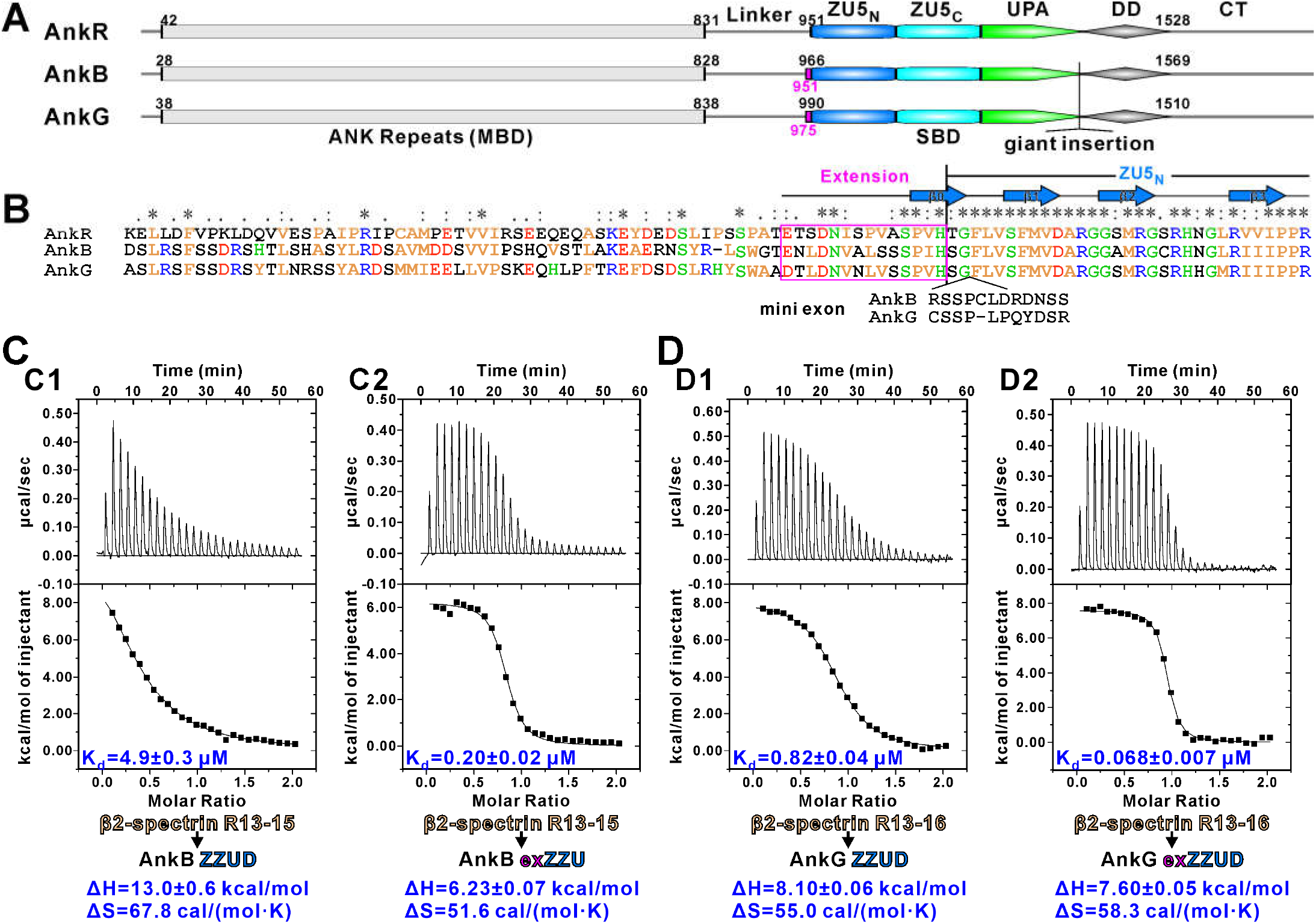
A conserved extension sequence is required for strong ankyrin/β-spectrin interactions. (A) Schematic diagram showing the domain organizations of AnkR/B/G. (B) Amino acid sequence alignment showing the conserved extension sequence N-terminal to ZU5_N_ among mouse AnkR, human AnkB and rat AnkG. “*”, “:”and “.” indicate fully conserved, highly conserved, and conserved residues, respectively. (C) ITC results showing that AnkB exZZU binding to β2-spectrin R13-15 is over 20 fold stronger than AnkB ZZUD without the extension. (D) ITC results showing that AnkG exZZUD binding to β2-spectrin R13-16 is over 10 fold stronger than AnkG ZZUD without the extension

A number of evidences imply that ankyrin and spectrin isoforms may have pairwise binding specificities at various subcellular microdomains. For example, AnkG specifically pairs with β2-spectrin in the lateral membranes of epithelial cells; AnkB forms complex with β2-spectrin throughout neuronal axons, whereas AnkG pairs specifically with β4-spectrin in AIS and NOR in axons [2]. Another interesting observation is that when AnkG was knocked out from axon, AnkR together with β1-spectrin instead of β4-spectrin became to be enriched in NOR [46]. Reciprocally, the β1-spectrin/AnkR pair instead of the β1-spectrin/AnkG complex became clustered in NOR after removal of β4-spectrin from neurons [46,47]. These observations suggest possible existence of specific pairings between isoforms of ankyrins and spectrins in living cells, though with totally unknown underlying molecular mechanisms.

Spectrins together with F-actin and some actin associated proteins like adducin form an elastic cytoskeleton network near the inner surface of plasma membranes. Ankyrins bind to integral membrane proteins (e.g. anion exchanger, L1CAM or E-cadherin) through their MBD and bind to β-spectrins through their SBD to connect plasma membranes to the elastic cytoskeleton thereby strengthening membrane lipid bilayers for mechanical stability. Interestingly, except for the ankyrin/spectrin interactions, the pairwise binding affinities between key components within this network are all quite strong with dissociation constants (Kd, or half maximal concentration) of a few tens nanomoles (Figure S1). For example, the half maximal concentration of the spectrin/actin interaction is ~15-75 nM [48], spectrin/adducin interaction is ~80-100 nM [49,50], ~60 nM for the adducin and F-actin tip interaction [48], and ~10 nM between α and β-spectrin [51]. Also, on the membrane-cytoskeleton interface, the Kd values of the ankyrin/anion exchanger complex is ~10 nM [1] and the ankyrin/neurofascin is ~50-500 nM [1,39]. These strong interactions presumably are fitting with the strong mechanostability roles of the ankyrin and spectrin/actin cytoskeleton assembly at the membrane microdomains. Curiously, the reported binding affinity between ankyrin and spectrin is ~6.8 μM [40], which is much weaker than the rest of the interactions in the network. Considering that ankyrin-spectrin interaction is the major linkage between the plasma membranes and the actin cytoskeleton of this network, the reported weak affinity between ankyrin and spectrin may not be able to support the stability of the network. Thus, we hypothesized that either the reported affinity on the ankyrin/spectrin interaction is incomplete or there are certain uncharacterized regulatory mechanisms for the ankyrin-spectrin interactions.

Although the ankyrin ZU5_N_-ZU5_C_-UPA tandem (ZZU) forms a compact supramodule [40], the bindings of ankyrins to spectrins seem to only require the first ZU5 domain (ZU5_N_) of the tandem [52,53]. Here, we discovered that an extension sequence preceding the ZU5_N_ of the ZZU tandem of ankyrins can dramatically enhance their spectrin binding affinities. We solved the crystal structures of AnkB exZZU in complex with repeats 13-15 (R13-15) of β4-spectrin and AnkG exZZU in complex with R13-15 of β2-spectrin and with R13-15 of β4-spectrin. These structures shed light on how the extension sequence in exZZU can enhance the interactions between ankyrins and spectrins. Our structures also reveal that, in addition to the ZU5_N_ domain, the UPA domain in the exZZU supramodule of ankyrins also directly contacts with spectrins. The biochemical and structural results presented in this study may serve as a foundation for understanding specific functional pairings between ankyrins and spectrins in living cells.

## Results

### A short extension N-terminal to ZU5_N_ enhances the ankyrin/spectrin interactions

Amino acid sequence analysis discovered that a short stretch of residues immediately preceding the ZU5_N_ domain of ankyrins are highly conserved during evolution and are very similar among AnkR/B/G (Figure 1A and 1B). Since the residues immediately following this extension (i.e. the β1 strand of ZU5_N_) are directly involved in binding to spectrins based on the structure of the AnkR ZU5_N_ in complex with R13-15 of β1-spectrin [52] (PDB ID: 3KBT; Figure 1B), we speculated that this extension may participate in ankyrins’ binding to spectrins. As in living cells, β2-spectrin can bind to both AnkB and AnkG [2], we therefore chose β2-spectrin as the representative β-spectrin to test our hypothesis. Interestingly, with this extension sequence (15 aa, boxed in pink in Figure 1A&B) appended to AnkB ZU5_N_-ZU5_C_-UPA (defined as exZZU, aa 951-1443), the binding affinity between β2-spectrin and AnkB exZZU (Kd ~0.20 μM) is >20 folds stronger than that between β2-spectrin and AnkB ZU5_N_-ZU5_C_-UPA-DD tandem (ZZUD) (aa 966-1535) (Kd values of ~0.20 μM *vs* ~4.9 μM; Figure 1C). Inclusion of additional N-terminal sequence as well as the death domain (i.e., an AnkB fragment containing aa 930-1535) did not further enhance the affinity of AnkB to β2-spectrin (Figure S2A), suggesting that this exZZU module is sufficient to achieve the strong binding between AnkB and β2-spectrin. Similarly, inclusion of this extension also enhanced AnkB’s binding to β4-spectrin (Figure S2B) and AnkG’s binding to β2-spectrin (Figure 1D). We have also investigated the role of the extension sequence in AnkB’s binding to β1-spectrin. Although AnkB is not known to pair with β1-spectrin in living cells, the extension sequence also enhanced the binding between AnkB and β1-spectrin by ~12 folds (Figure S2C). Taken together, the above biochemical studies indicated that the extension sequence N-terminal to the ZU5_N_ can enhance pairwise bindings between ankyrins and β-spectrins. Importantly, the Kd values between ankyrins and β-spectrins are in the range of tens to hundreds of nanomoles, matching with the rest of the strong pairwise interactions in the ankyrin/actin cytoskeleton networks shown in Figure S1. Our results further suggested that the interactions between various combinations of ankyrin and β-spectrin pairs all appear to be strong interactions (i.e. there appears no pairwise specificity code between specific pairs of ankyrins and β-spectrins observed in cells).

### Crystal structure of the β2-spectrin/AnkG exZZU complex

To elucidate detailed mechanisms underlying the N-terminal extension-mediated enhancement of interactions between ankyrins and β-spectrins, we first crystalized β2-spectrin repeat 13-15 in complex with AnkG exZZU (aa 975-1465) and solved the complex crystal structure at the resolution of 3.3 Å (Table 1). Consistent with the apoform structure of the AnkB ZU5_N_-ZU5_C_-UPA tandem [40], the AnkG ZU5_N_-ZU5_C_-UPA tandem also forms a supramodule with a cloverleaf-like architecture (Figure 2A), suggesting the formation of the ZZU supramodule is a common feature shared by all three ankyrin family members. The R13-R15 repeats of β2-spectrin contacts with the surfaces of ZU5_N_ and UPA. The interface between β2-spectrin R14 and AnkG ZU5_N_ is nearly identical to that between β1-spectrin R14 and AnkR ZU5_N_ as shown in the structure of β1-spectrin R13-15 in complex with the solo ZU5_N_ of AnkR (Figure S3) [52]. Although the ZZU supramodule provides a larger spectrin binding interface, R13 of β2-spectrin is not involved in the AnkG binding. This is consistent with our biochemical result showing that deleting R13 does not impair ankyrin/β-spectrin interaction (Figure S2A).

**Figure 2:**
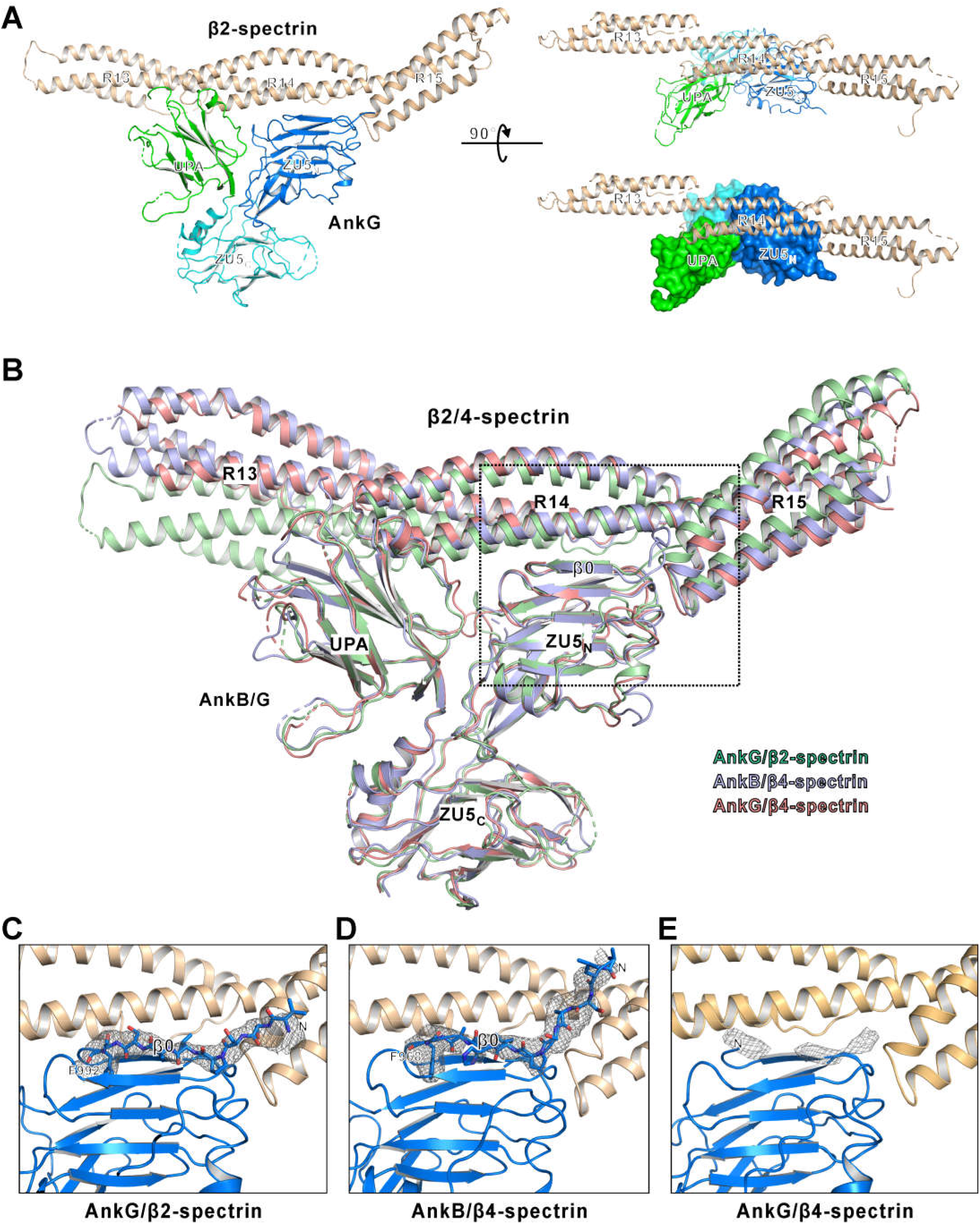
Overall structure of AnkG exZZU/β2-spectrin R13-15 complex and common features revealed by the three complex structures. (A) Ribbons diagram representation (*left & right top*) and combined ribbon and surface representations *(right bottom)* of the crystal structure of AnkG exZZU/β2-spectrin R13-15. ZU5_N_ (blue), ZU5_C_ (cyan), UPA (green), and β2-spectrin (gold) are drawn in their specific colors and this color code is used throughout the manuscript except as otherwise indicated. (B) Superposition of all the three structures showing that the binding modes are very similar. In this superposition, ZU5_N_ was used as the reference for the structural alignment. (C-E) Omit maps showing the formation of a β strand by the extension in AnkG/β2-spectrin (C), AnkB/β4-spectrin (D), and AnkG/β4-spectrin (E). Each Fo-Fc density map was generated by deleting the extension sequence from the final model and contoured at 2.5 σ.

**Table 1:**
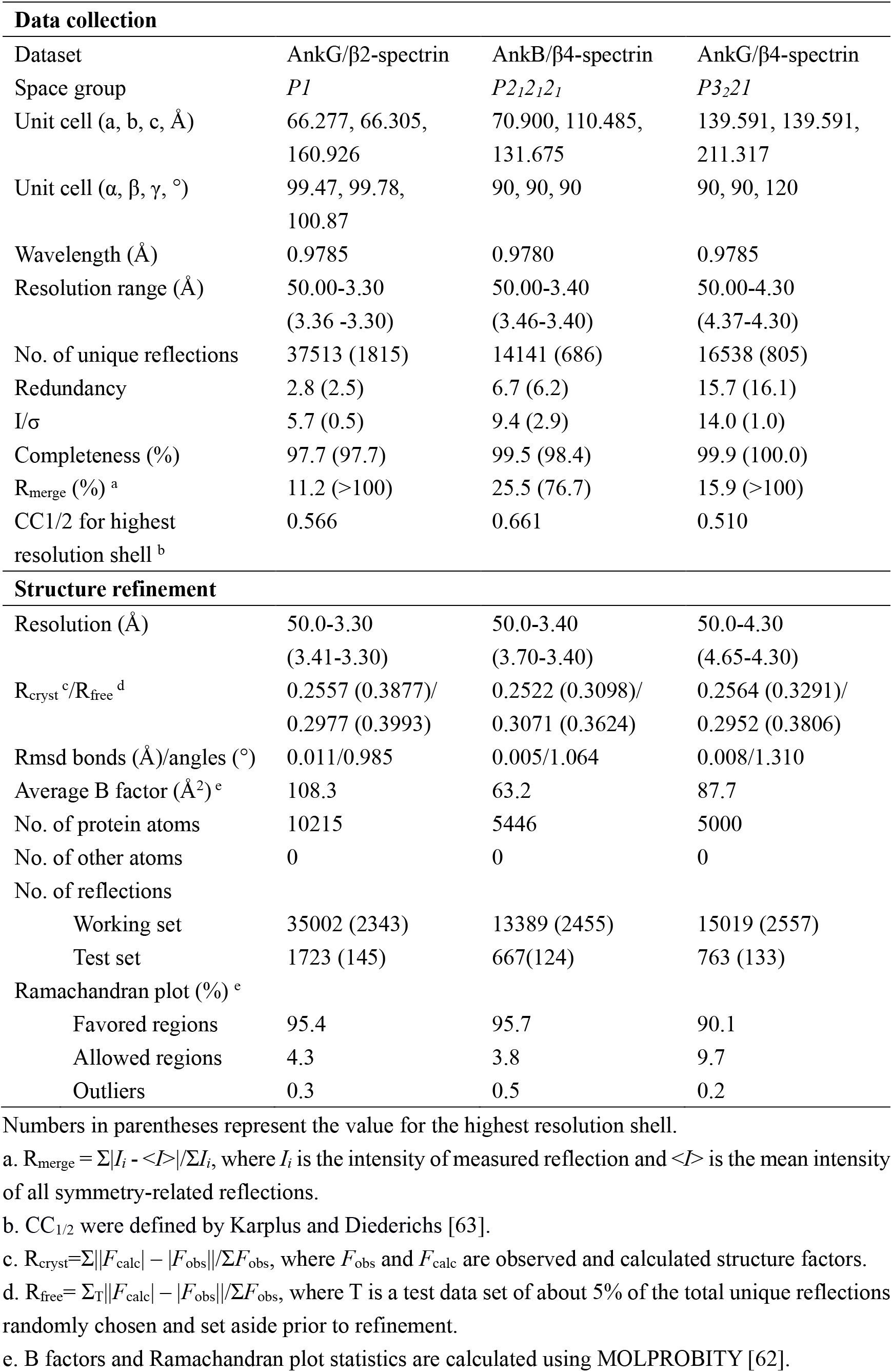
Statistics of X-ray Crystallographic Data Collection and Model refinement

Guided by the AnkG/β2-spectrin complex structure, we were able to optimize construct boundaries of the AnkB/β4-spectrin and AnkG/β4-spectrin complexes for crystallizations and solved the structures of these two complexes at resolutions of 3.4 Å and 4.3 Å, respectively (Figure 2B-E and Table 1). The overall architectures of the three pairs of ankyrin exZZU/β-spectrin R13-15 structures are essentially the same, including the cloverleaf-like ZU5_N_-ZU5_C_-UPA supramodule and the main ZU5_N_/β-spectrin R14 interface. The orientations of R13 and R15 with respect to R14 in three structures are slightly different (Figure 2B), which is not surprising given that the conformations between spectrin repeats are known to be partially elastic in spectrins [41–44].

### Structural basis for the enhanced spectrin binding by the N-terminal extension of ankyrins

Based on the three complex structures, we analyzed how the N-extension can enhance ankyrin’s binding to β-spectrin. When building the structure models, we observed clear electron densities N-terminal to F992 in AnkG or F968 in AnkB (Figure 2C and 3D). These electron densities correspond to the residues from the N-terminal extension. For the AnkG/β2-spectrin and AnkB/β4-spectrin structures, an additional β strand was modeled and denoted as β0 corresponding to the N-terminal extension for each ankyrin exZZU (Figure 2C and 2D). The low resolution of the AnkG/β4-spectrin structure did not allow us to confidently build the atomic model, but obvious electron densities in the difference map also indicate the formation of β0 in the exZU5_N_ in AnkG when in complex with β4-spectrin (Figure 2E).

**Figure 3:**
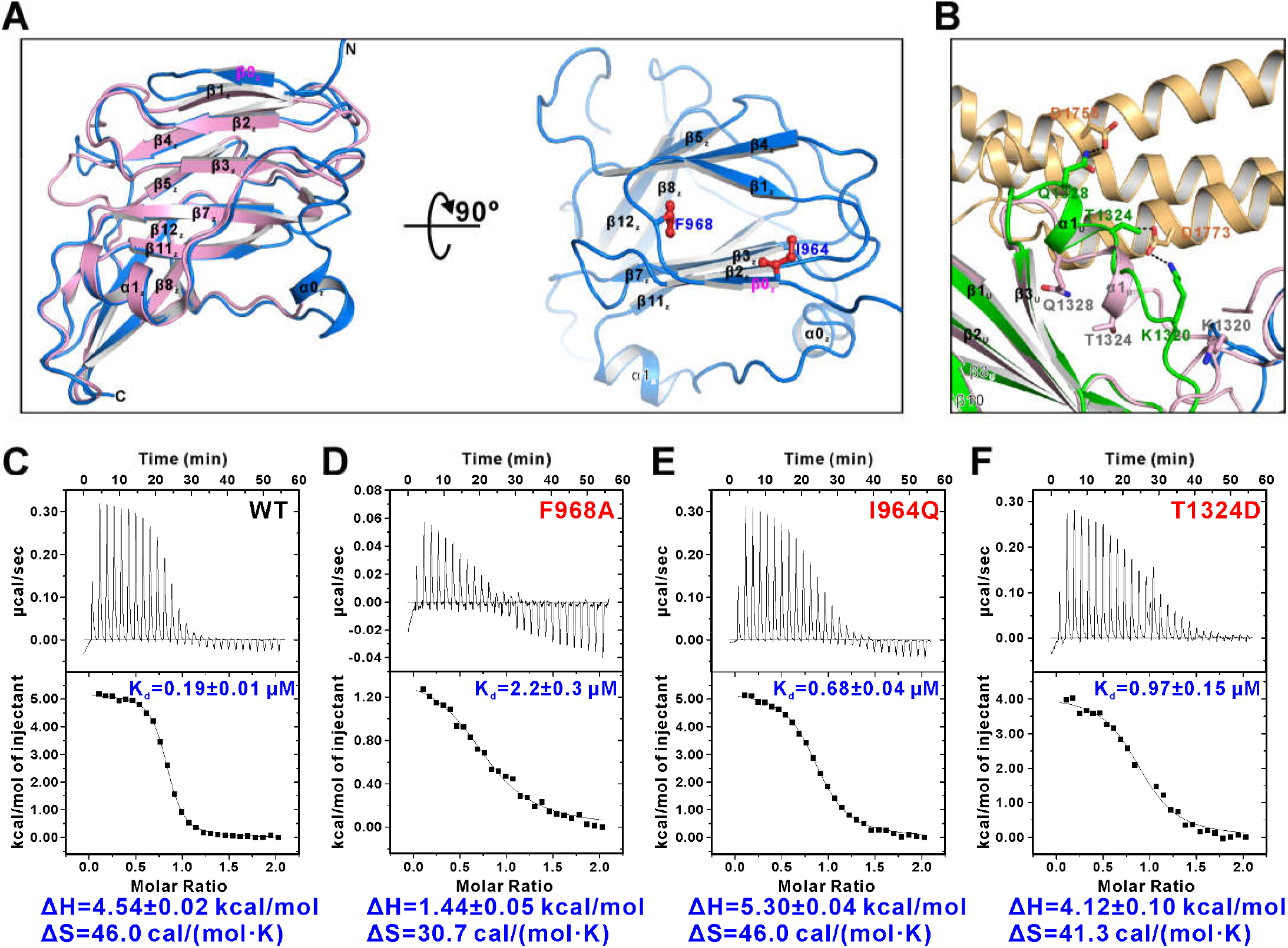
Detailed interaction of the ankyrin/β-spectrin complexes using the AnkB/β4-spectrin complex as an example. (A) The additional β0 formed by the extension packs to the ZU5_N_ domain mainly through hydrophobic interactions. The ZU5_N_ of the apo AnkB ZZUD structure is colored in pink. (B) UPA domain is involved in the interaction by hydrogen bonding. The UPA domain of the apo AnkB ZZUD structure is colored in pink. (C-F) ITC results showing that mutating the key residues in the interface between AnkB exZZU and β4-spectrin weakened the binding.

Surprisingly, there is no direct contact between β-spectrin and ZU5_N_ β0 in both AnkG/β2-spectrin and AnkB/β4-spectrin complex structures. We hypothesized that the enhanced binding might be due to the stabilization of ZU5_N_ by the additional β strand (i.e. β0) formed by the N-terminal extension. The β0 strand is anti-parallelly paired with β1, making ZU5_N_ a 13-stranded β-barrel (Figure 3A). Two hydrophobic residues in the extension sequence (I964 and F968 in AnkB exZZU, V986 and F992 in AnkG exZZU; Figure 1B) insert into the hydrophobic core of ZU5_N_, which should stabilize the folding of the ZU5 domain. Nonetheless, these two residues are not absolutely required for the overall folding of the remaining 12 β strands of ZU5_N_ as I964 is missing and side-chain of F968 is facing outside (i.e. exposed to solvent) in the previous solved apo-AnkB ZZUD structure (PDB ID: 4D8O). Substitution of F968 by alanine decreased the AnkB exZZU/β2-spectrin binding by more than ten folds (Figure 3D). Similarly, replacing I964 by a polar glutamine also weakened the binding by about four-fold (Figure 3E).

The above biochemical and structural analysis suggested that, by forming an additional β strand, the N-terminal extension can stabilize the ZU5_N_ structure and consequently enhance the bindings of ankyrin exZZU to β2-spectrin.

### The UPA domain is also involved in the ankyrin-spectrin interaction

Another new finding from the three complex structures solved in this study is that the UPA domain of ankyrin exZZU directly participates in binding to β-spectrin. By superimposing the UPA domains’ structures from the AnkB exZZU/β4-spectrin complex and from the apo-AnkB ZZU structure (Figure 3B), we could observe an obvious shift and rotation of the α1 helix. Such shift and rotation facilitate a few direct polar interaction between residues from α1 of UPA and R14 of β-spectrin. For example, T1324 and Q1328 on AnkB exZZUD form hydrogen bonds with D1773 and D1755 on β4-spectrin repeat 14, respectively. K1320 in the loop preceding α1 of UPA also forms a salt bridge with D1773 from β4-spectrin. Consistent with the above structural analysis, substitution of T1324 in α1 of UPA with Asp reduced the interaction between AnkB exZZU and β4-spectrin by about five-fold (Figure 3F).

### The exZZU supramodule of ankyrins functions as an integral unit for binding to β-spectrin

The finding that both the ZU5_N_ and UPA domains of ankyrins are directly involved in binding to β-spectrins prompted us to propose that the overall conformation (i.e. the tertiary structure) of the exZZU supramodule is important for ankyrin-spectrin interactions (Figure 4A). Our previous [40] and currently solved structures revealed that R1029 from AnkB ZU5_N_ is critical for the ZU5_N_/UPA coupling by forming salt bridges with a negatively charged residue (D1319 as revealed by the previous AnkB ZZUD structure or D1318 as revealed by the structure solved here) from UPA domain (Figure 4B). The residues corresponding to R1029 and D1318/1319 are completely conserved in all isoforms of ankyrins throughout the evolution (see Figure 2 in Wang et al., 2012 for detailed sequence alignment analysis). We replaced R1029 of AnkB with Glu to destabilize the ZU5_N_/UPA coupling in the exZZU supramodule. The R1029E mutation reduced AnkB exZZU’s binding to β2-spectrin by ~5 folds (Figure 4C and 4D). The above biochemical and structural analyses suggest that the ZU5_N_/UPA coupling stabilizes the tertiary structure of the ankyrin exZZU supramodule, which is required for the high affinity interactions between ankyrins and β-spectrins.

**Figure 4:**
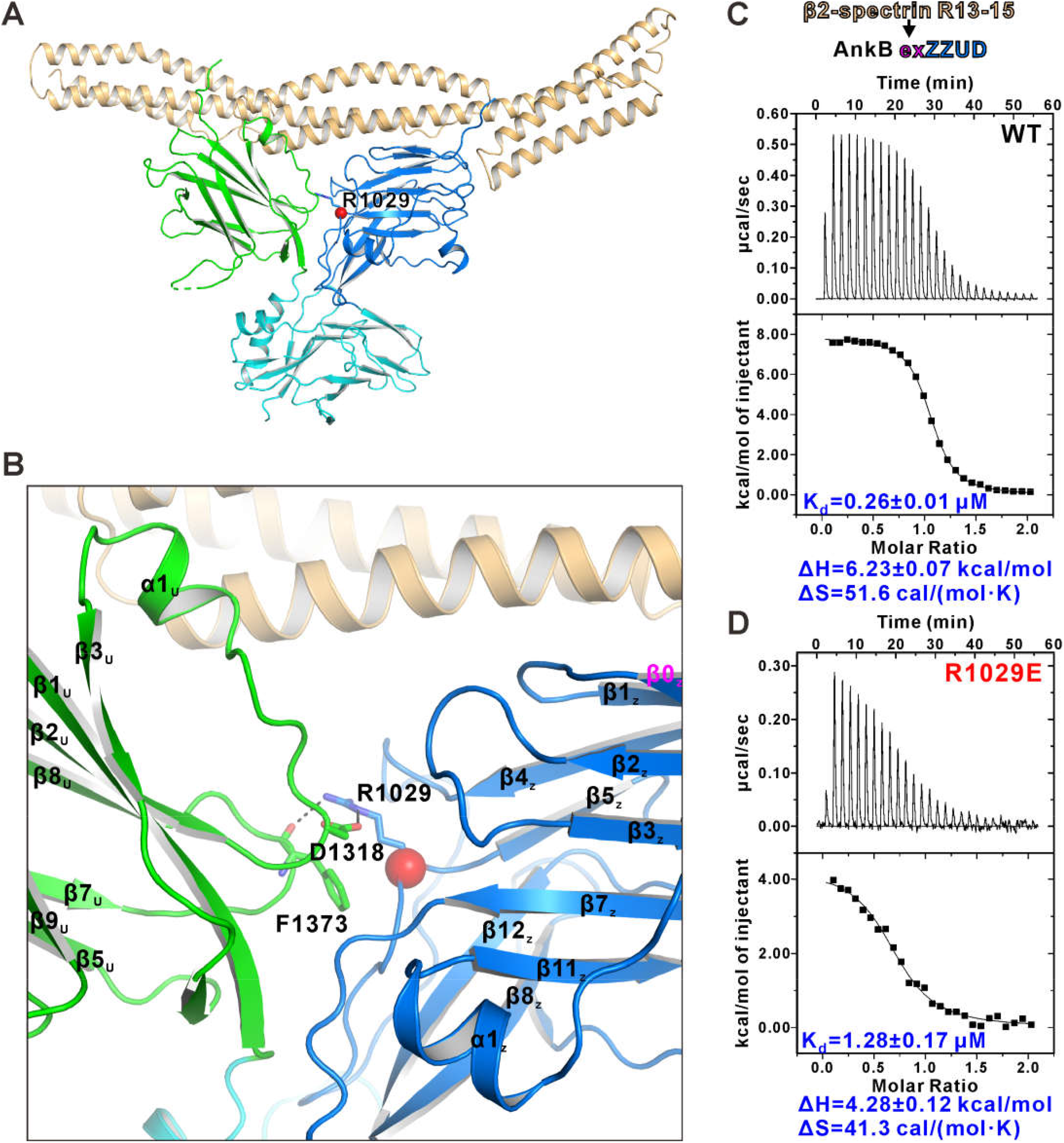
ZZU supramodule as an integral unit to bind to β-spectrin. (A) R1029 (the red sphere) is located in the ZU5_N_/UPA interface. (B) R1029 is critical for the ZU5_N_/UPA coupling by forming a salt bridge with D1318. (C-D) A charge reversal mutation of R1029 decreased the binding between AnkB exZZU and β2-spectrin.

## Discussion

In this study, we identified a short and evolutionally conserved sequence immediately preceding the ZU5_N_ domain of AnkB and AnkG. It can significantly enhance the binding between AnkB/AnkG and β-spectrins. We demonstrated that the binding affinities between AnkB/AnkG and β-spectrins are in similar range as those very strong pairwise bindings between the ankyrin/actin cytoskeleton network components, which are critical for stable membrane micro-domain assemblies (Figure S1; [1,39,48–51]). The structures of AnkB/AnkG exZZU in complex with R13-15 of β2/β4-spectrins illustrate that the N-terminal extension stabilizes the ZU5_N_ domain structure by forming an additional β strand and thus enhances β-spectrin binding. The structures of the ankyrin/spectrin complexes solved in this study further reveal that the UPA domain of ankyrins directly participates in binding to β-spectrin. We provided biochemical evidence to show that the tertiary structural arrangement of the exZZU supramodule is important for the interactions between ankyrins and β-spectrins, implying that long-range conformational perturbations may regulate ankyrin/β-spectrin interactions. This tertiary structural requirement of ankyrin exZZU for binding to β-spectrin may provide a possible explanation to why certain disease mutations in the exZZU supramodule may alter ankyrin’s binding to β-spectrin, even if such mutation sites are away from the β-spectrin binding surface.

A splicing switch was recently reported to modulate the AnkG/β4-spectrin interaction in AIS during neuronal development [54–56]. The splicing factor Rbfox or Elavl3 can regulate the insertion of a mini-exon coding extra 11 amino acid residues into the site between the extension sequence and the canonical ZU5_N_ domain in both AnkG and AnkB (see Figure 1B for the insertion site). For AnkG splicing isoform containing this mini-exon, the binding of AnkG to β2- or β4-spectrin is largely decreased as demonstrated by immunoprecipitation experiments [54]. This result is in line with our structural and biochemical data presented in this study. The insertion of the 11-residue fragment encoded by the mini-exon is expected to disrupt the β0 strand formed by the N-terminal extension sequence in AnkG (and in AnkB and AnkR as well) and thereby decrease the binding affinity between AnkG and β2-/β4-spectrin. Thus, in addition to directly enhancing the interactions between ankyrins and spectrins, the N-terminal extension of the ankyrin exZZU supramodule also serves as a splicing-mediated switch for modulating their interactions.

A surprising and unexpected finding from the current structural and biochemical study is that AnkB and AnkG use very similar binding mode to interact with different β-spectrins and with comparable affinities (Figure 1 and Figure S2). This finding suggests that the specific pairings between isoforms of ankyrins and β-spectrins observed in distinct membrane microdomains of living cells are not solely determined by the high affinity interactions between the exZZU of ankyrins and R13-15 of β-spectrins. There should be certain additional mechanism(s) that can modulate specific pairings between ankyrins and β-spectrins. Supporting this speculation, a recent study showed that the giant insertion encoded by a single exon coding >3000 amino acid residues in 480-kDa AnkG is essential for the recruitment of β4-spectrin to AIS [57]. Further studies will be required to understand the detailed mechanisms underlying the specific pairings between ankyrins and β-spectrins.

## Materials and methods

### Constructs, protein expression and purification

All of the AnkB constructs coding sequences were PCR amplified from the full-length human 220-kDa AnkB cDNA plasmid (residue numbers referred to NP_001341186.1). All of the AnkG constructs were originated from a rat 270-kDa cDNA (residue numbers according to GenBank ID: AAC78143.1). Both of the AnkB and AnkG template plasmids are gifts from Dr. Vann Bennett at Duke University. β-spectrins’ coding sequences were PCR amplified from mouse brain or muscle cDNA libraries. The corresponding residue numbers of different spectrin repeats are summarized in Table 2. All point mutations were generated by the Quick Change site-directed mutagenesis kit (Agilent, CA) and confirmed by DNA sequencing. All of these coding sequences were cloned into home-modified pET32a vectors for protein expression.

**Table 2:** Summary of β-spectrin constructs’ boundaries

| **Species** | **Spectrin** | **Spectrin Repeats** | **Residue Number** |
| --- | --- | --- | --- |
| mouse | β1-spectrin<br>NP_038703.3 | 13-16 | 1583-2010 |
| mouse | β2-spectrin<br>NP_787030.2 | 13-15 | 1591-1904 |
|  |  | 13-16 | 1591-2018 |
|  |  | 14-15 | 1694-1904 |
| mouse | β4-spectrin<br>AAK38731.1 | 13-15 | 1611-1932 |
|  |  | 13-16 | 1611-2040 |
|  |  | 14-15 | 1714-1932 |

Recombinant proteins were purified as previously described [39]. In brief, 200 ng plasmids were transformed into 50 μL *Escherichia coli*. BL21 (DE3) (New England Biolabs) competent cells. On the second day, the cells were inoculated to 1-2 L Lysogeny broth medium and incubated at 37 °C with shaking at 200 rpm. When UV absorbance at 600 nm of the cultured cells reached 0.8, the cells were induced by adding 0.25 mM IPTG and the culture was shifted to 16 °C for 20 hours with shaking at 200 rpm. The cells were then pelleted by centrifugation at 3,000 g for 15 minutes, resuspended with 40 mL resuspension buffer (50 mM Tris, 500 mM NaCl, 5 mM imidazole, pH 8.0), and lysed by a high pressure homogenizer machine at 4°C, followed by high speed centrifugation at 39,000 g for 20 minutes. The supernatant was injected into a Ni^2+^-NTA agarose affinity column (containing 5 mL Ni^2+^-Sepharose 6 FF beads, GE Healthcare, Cat. 17531803). The column was incubated for 15 minutes and washed twice with 30 mL washing buffer (50 mM Tris, 500 mM NaCl, 15 mM imidazole, pH 8.0). Then the proteins on the column were eluted by 15 mL elution buffer (50 mM Tris, 500 mM NaCl, 1 M imidazole, pH 8.0). The eluted proteins were applied to a size-exclusion column (HiLoad 26/600 Superdex 200 pg column from GE Healthcare) with the buffer containing 50 mM Tris-HCl, 1 mM DTT, and 1 mM EDTA, pH 7.8 with 100 mM NaCl. The fused Trx-His_6_ tag was cleaved by incubating each recombinant protein with HRV 3C protease overnight at 4 °C. The cleaved tag of each protein was removed by another step of the size exclusion chromatography.

### Isothermal Titration Calorimetry (ITC) assay

Isothermal titration calorimetry (ITC) measurements were performed on a VP-ITC Microcal calorimeter (MicroCal, Northampton, MA) at 25 °C. All purified protein samples were in reaction buffer containing 50 mM Tris, 100 mM NaCl, 1 mM EDTA and 1 mM DTT at pH 7.5. Spectrin proteins in the concentrations of 100-200 μM were loaded into the syringe and ankyrin proteins in the concentrations of 10-20 μM were loaded into the sample cell of the machine. Each titration point was performed by injecting a 10 μL aliquot (the first titration point is 5 μL) of syringe protein into corresponding ankyrin protein samples in the cell at a time interval of 120 seconds. The titration data were analyzed using the program Origin7.0 and fitted by the one-site binding model.

### Crystallography

All crystals were obtained by hanging drop or sitting drop vapor diffusion methods at 16°C. Crystals of AnkG exZZU/β2-spectrin R13-R15 were grown in solution containing 10% w/v polyethylene glycol 6,000, 5% v/v (+/-)-2-methyl-2,4-pentanediol and 0.1 M HEPES pH 7.5. Crystals of AnkB exZZU/β4-spectrin R13-R15 were grown in solution containing 25% w/v pentaerythritol propoxylate 629 (17/8 PO/OH), 50 mM magnesium chloride and 0.1 M HEPES, pH 7.5. Crystals of AnkG exZZU/β4-spectrin R13-R15 were grown in solution containing 0.8 M potassium sodium tartrate tetrahydrate, 0.1 M Tris, pH 8.5 and 0.5% w/v polyethylene glycol monomethyl ether 5,000. Crystals were soaked in crystallization solution containing additional 20% glycerol for cryoprotection. All datasets were collected at the Shanghai Synchrotron Radiation Facility (BL17U1 or BL19U1) at 100 K. Data were processed and scaled using HKL2000 or HKL3000 [58].

Structures were solved by molecular replacement using PHASER [59] with AnkB ZZU (PDB: 4D8O) and β1-spectrin R13-R15 (PDB: 3KBT) as the searching models. Further manual model adjustment and refinement were completed iteratively using COOT [60] and PHENIX [61]. The final models were validated by MolProbity [63] and statistics are summarized in Table 1. All structure figures were prepared by PyMOL (http://www.pymol.org). The coordinates of the structures reported in this work are being deposited to PDB.

## Acknowledgements

We thank the Shanghai Synchrotron Radiation Facility (SSRF) BL19U1 and BL17U1 for X-ray beam time. This work was supported by grants from RGC of Hong Kong (16100517 and 16104518) and from Natural Science Foundation of Guangdong Province (2016A030312016) and Shenzhen Basic Research Grant (JCYJ20160229153100269). M.Z. is a Kerry Holdings Professor in Science and a Senior Fellow of IAS at HKUST.

## Supplemental Figure Legend

**Figure S1:**
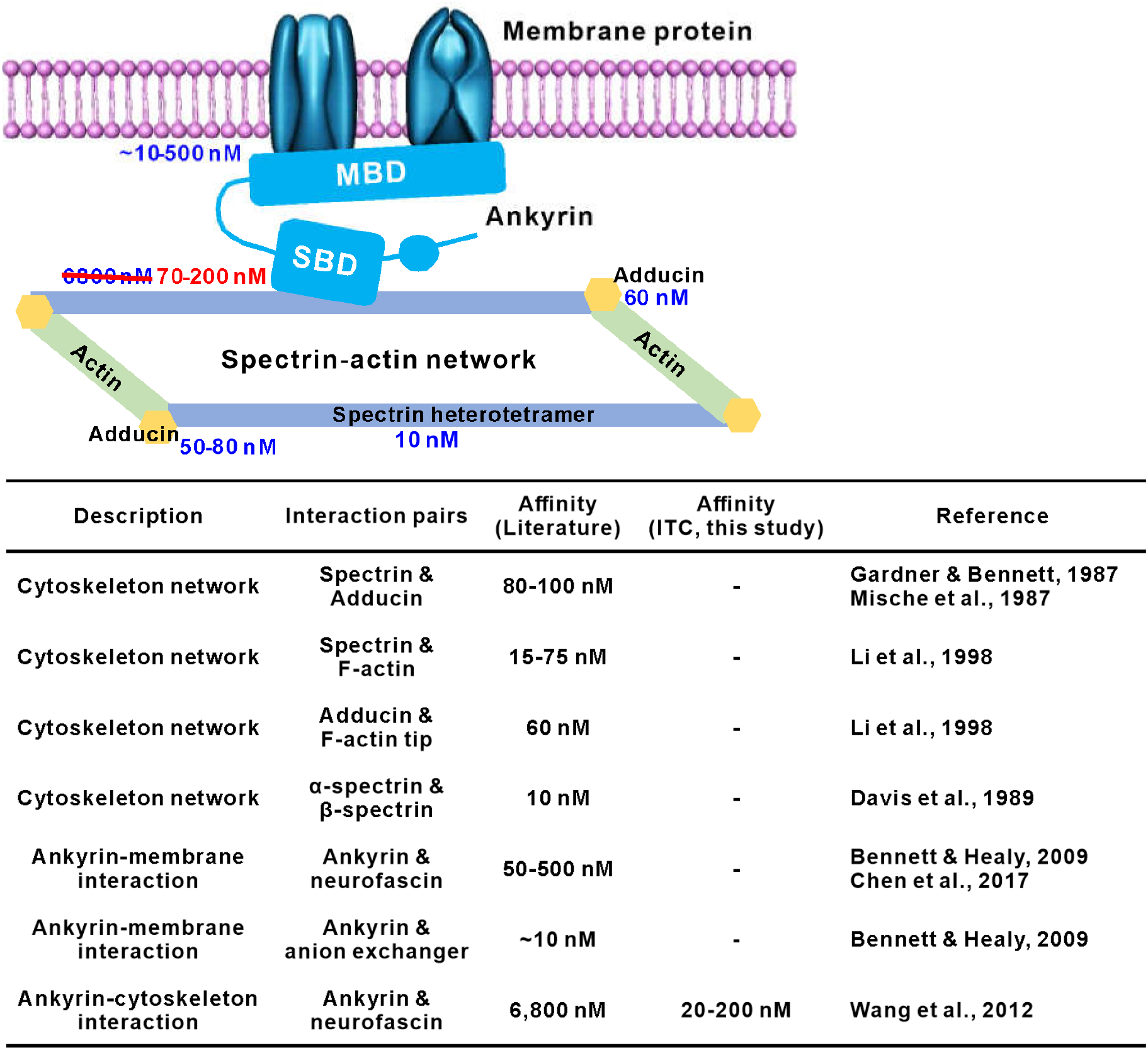
Schematics showing the layered arrangement of the membrane-ankyrin-spectrin network and affinities of interactions between key components. Ankyrin with its MBD, SBD and DD is shown in light blue; αβ-spectrin heterotetramers are represented by light purple columns; adducin is shown in yellow and actin is shown in light green. Table in the bottom shows literature reported binding affinities (dissociation constants or half maximal concentrations) between key components. *“* -” means not determined in this study.

**Figure S2:**
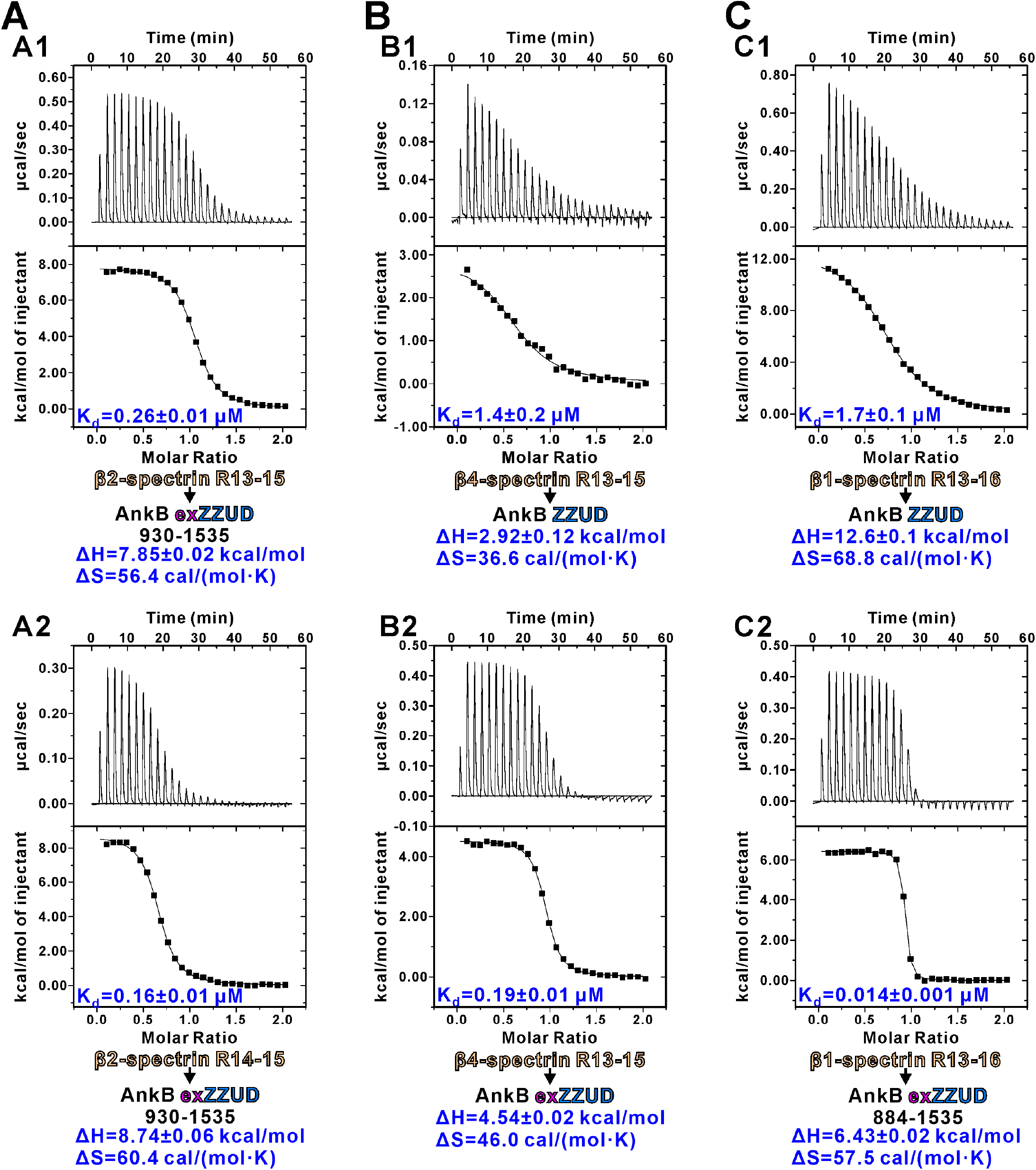
ITC data showing different AnkB constructs binding to β2-(A) or β4-(B) or β1-spectrin (C).

**Figure S3:**
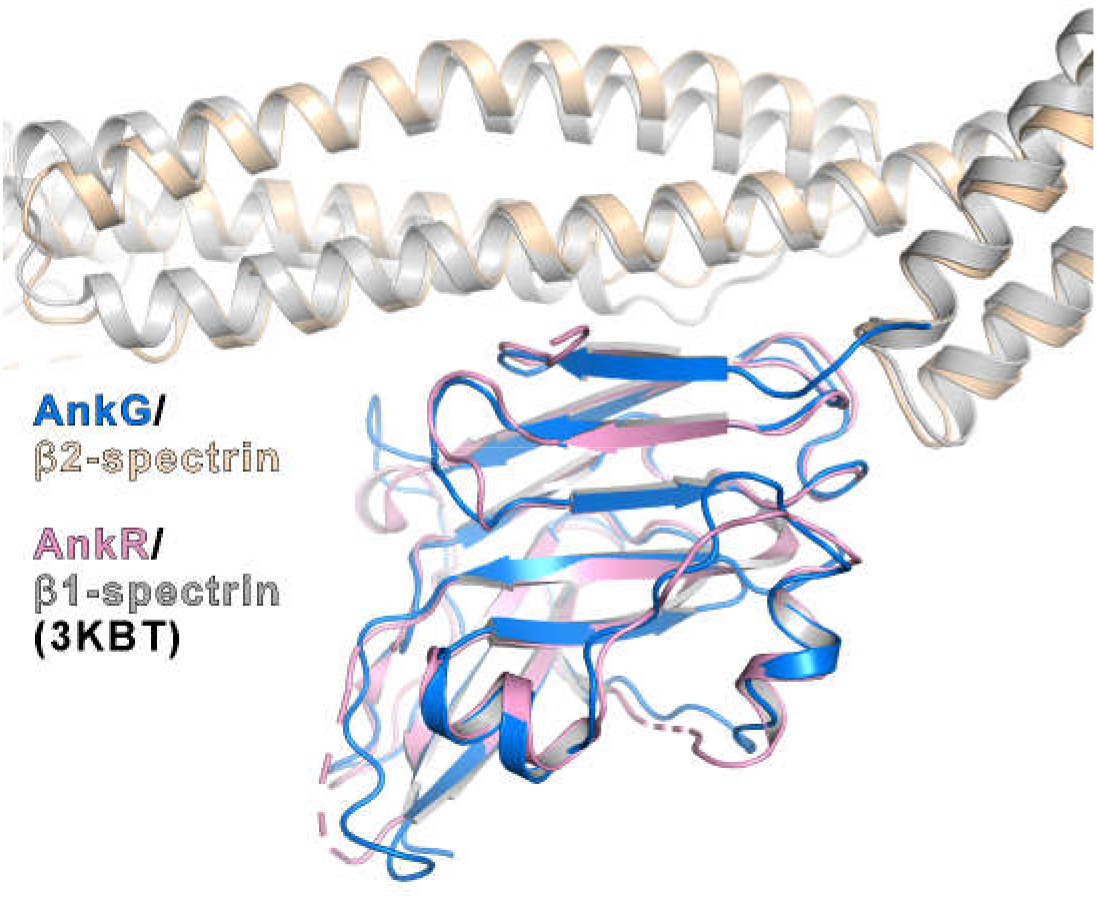
Structure alignment of the β1-spectrin/AnkR complex and the β2-spectrin/AnkG complex. The AnkR ZU5_N_ domain is shown in pink and β1-spectrin is shown in grey (PDB ID: 3KBT). The AnkG ZU5_N_ domain is shown in blue and β2-spectrin shown in gold. For simplicity, the ZU5_C_ and UPA domains are not drawn.

